# MorphCell learns interpretable shape representations of 3D cellular morphology

**DOI:** 10.64898/2026.09.17.752374

**Authors:** Guowei Zhang, Guan Wang, Runzhou Cao, Jiansheng Guo, Shan Xu, Zeyu Yu, Yiyan Zheng, Yong He, Xuping Feng

**Affiliations:** College of Biosystems Engineering and Food Science, Zhejiang University, Hangzhou, China; School of Artificial Intelligence and Information Engineering, Zhejiang University of Science and Technology, Hangzhou, China; Center of Cryo-Electron Microscopy, Zhejiang University School of Medicine, Hangzhou, China; Westlake Laboratory of Life Sciences and Biomedicine, Hangzhou, China; The Rural Development Academy & Agricultural Experiment Station, Zhejiang University, Hangzhou, China

## Abstract

Three-dimensional (3D) cell morphology provides a measurable phenotype of cellular state and function, yet learning transferable and interpretable representations across biological systems and imaging modalities remains challenging. Here we introduce MorphCell, a self-supervised framework for learning 3D cellular morphology representations from cell-surface point clouds. MorphCell uses cross-view reconstruction and spherical self-reconstruction to capture global and local geometric features and reconstruct complete 3D surfaces. Pretrained on diverse non-biological 3D geometries, MorphCell transfers to biological datasets without task-specific fine-tuning, providing a common representation across imaging modalities and biological contexts. We demonstrate this transferability across confocal microscopy, X-ray microscopy and volume electron microscopy, spanning red blood cells, wheat root cells and colon epithelial cells. MorphCell captured biologically relevant 3D shape variation and outperformed baseline representations in multiple classification tasks. Separating physical scale from the learned shape representation further revealed context-dependent contributions of shape and scale: shape alone outperformed shape-scale fusion for red blood cell morphotype classification, whereas combining shape with the nuclear-to-cytoplasmic volume ratio increased balanced accuracy for cancerous versus non-cancerous colon cells from 62.7% to 92.8%. MorphCell therefore provides a general framework for representing, reconstructing and interpreting 3D morphology, enabling the discovery of biologically meaningful morphological features beyond conventional measurements of scale.

## Main

Cellular morphology provides a measurable phenotype associated with cell state and function^1,2^. During embryogenesis and tissue morphogenesis, changes in three-dimensional (3D) cell shape and volume are coordinated with lineage specification, cell-fate transitions and the establishment of tissue architecture^3,4^. Beyond reflecting cell state, specific aspects of morphology can also be associated with biological processes. Interphase morphology can affect the mode, symmetry and outcome of mitosis^5^, while cell size varies with G1/S transition in mammalian stem cells in vivo^6^. However, conventional morphological profiling commonly relies on two-dimensional (2D) images, which cannot fully capture complex 3D cell shapes^7–10^. Moreover, existing 3D morphological analyses often emphasize size-related descriptors, such as cell volume, surface area and length, while overlooking complex shape variations, including local protrusions, depressions and other surface irregularities. These morphological features may contain biologically relevant information that is not captured by size measurements alone.

Advances in light-sheet microscopy, X-ray microscopy (XRM) and volume electron microscopy (vEM) now enable cellular morphology to be resolved in three dimensions across diverse biological specimens^11–13^. Cell surfaces segmented from these volumetric images can be represented as meshes or point clouds, providing a common geometric description of complex 3D morphology^14–16^. These geometric representations provide a basis for quantifying biologically relevant variation in 3D shape. However, few existing approaches both quantify such variation and preserve an interpretable connection to the underlying 3D geometry.

Methods for quantitative morphology analysis have traditionally relied on biophysical descriptors or spectral models. While biophysical descriptors yield directly interpretable geometric measurements, they are constrained by human-selected features^14,17^. By contrast, spherical-harmonic models provide reconstructable representations, yet struggle to capture irregular geometries and fine local detail due to their dependence on surface parameterization and a finite expansion order^18–20^. To reduce reliance on predefined features, learned image models have increasingly been adopted for unsupervised profiling^21,22^. However, because most architectures operate on 2D images, they fail to capture the complete surface geometry of 3D cells. Although recent point-cloud models extend representation learning into 3D space^15,16^, combining transferability across biological datasets with biological interpretability remains a challenge.

Here, we present MorphCell, a self-supervised framework for learning transferable and reconstructable representations of 3D cell-surface morphology. We evaluate MorphCell across datasets of red blood cells, wheat root cells and colon epithelial cells acquired by confocal microscopy, XRM and vEM, respectively. We examine which aspects of 3D morphology are captured by the learned representations and evaluate their transferability through cell classification tasks without biological task-specific training. These tasks include distinguishing red blood cell morphotypes, classifying wheat root cells by axial developmental zone and radial tissue region, and distinguishing cancerous from non-cancerous colon epithelial cells. We further investigate how the relative contributions of physical scale and 3D shape to cell classification vary across these biological tasks.

## Results

### MorphCell learns reconstructable representations of 3D cell morphology

We developed MorphCell, a representation-learning framework for 3D cell-surface point clouds. MorphCell combines cross-view reconstruction with spherical self-reconstruction. Cross-view reconstruction promotes the learning of robust morphology representations, whereas self-reconstruction recovers the complete 3D morphology from each representation.

3D microscopy stacks were converted into surface meshes and sampled as point clouds (Fig. 1a; see Methods). Principal-axis alignment standardized their orientation before encoding. MorphCell then transformed each point cloud into a morphology representation for 3D cell reconstruction, cell profiling and morphotype classification.

**Fig. 1.**
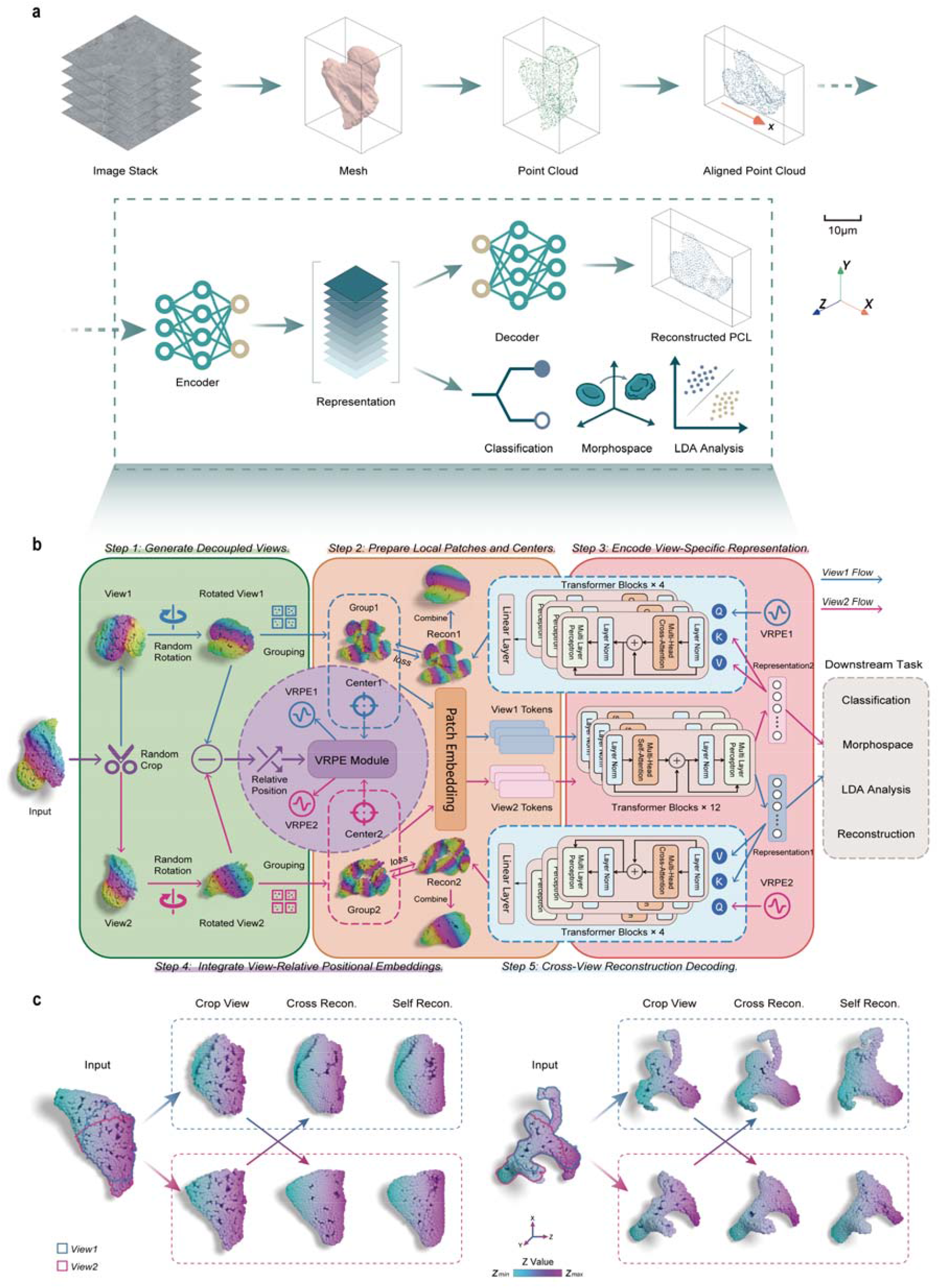
MorphCell learns reconstructable representations of 3D morphology. **a**, Overview of the MorphCell workflow. Raw 3D microscopy stacks are converted into surface meshes, sampled as point clouds and aligned along their principal axes. MorphCell encodes each aligned point cloud into a morphology representation for reconstruction and downstream analysis. Scale bar, 10 μm. **b**, Cross-view reconstruction framework. Each point cloud is randomly cropped into two local views, which are independently rotated and grouped into local patches. A shared encoder extracts a representation from each view. View-relative positional embeddings (VRPEs) encode the relative positions of the two views and guide reconstruction of the opposite view. **c**, Cross-view and self-reconstruction examples for a colon epithelial cell and an intracranial vessel segment^24^. Cross-view reconstruction recovers the opposite cropped view, whereas self-reconstruction recovers the complete point cloud from each view-specific representation. Blue and magenta boxes indicate the two view directions. Points are colored by their z coordinate.

Cross-view reconstruction served as the primary pretraining strategy^23^ (Fig. 1b; see Methods). Independent cropping and rotation generated two distinct local views of each point cloud. A shared encoder extracted a representation from one view, which was combined with view-relative positional information to reconstruct the other view. The model therefore learned geometric relationships between different surface regions rather than simply reproducing its input. This more demanding pretraining task promoted the learning of informative morphology representations.

Cross-view reconstruction alone could not recover a complete point cloud from a single representation. We therefore developed a Spherical-Query Transformer Decoder (SQTD) for self-reconstruction (Supplementary Fig. 1; see Methods). SQTD used uniformly sampled points on a Fibonacci sphere as an initial template and predicted a displacement for each point, progressively deforming the template into the target morphology. This reconstruction branch enabled visualization of the 3D geometry encoded by each representation.

We qualitatively evaluated both reconstruction branches using biological samples excluded from pretraining (Fig. 1c). Cross-view reconstruction recovered the geometry of the opposite local view in both reconstruction directions. Self-reconstruction recovered a complete point cloud from each view-specific representation. For vEM data from colon epithelial cells, the reconstructions retained the principal surface contours and local undulations. For intracranial vasculature acquired by time-of-flight magnetic resonance angiography (TOF-MRA)^24^, they retained the major branches of the tubular structures. These results show that MorphCell learned robust 3D morphology representations for subsequent cell profiling and morphotype classification, while retaining self-reconstruction capability for morphological interpretation.

### Shape-driven representations distinguish RBC morphotypes without fine-tuning

To evaluate transferability without biological task-specific training, we applied the pretrained MorphCell to a 3D red blood cell (RBC) dataset acquired by confocal laser scanning microscopy^25^. The dataset comprised seven characteristic RBC morphotypes, including discocyte, stomatocyte, spherocyte, knizocyte and echinocyte I– III (Fig. 2a). We first examined physical scale (defined as maximum cell diameter) of these morphotypes (Fig. 2b). Physical scale varied among morphotypes, with spherocytes showing the smallest average values. However, its broad within-class variability and substantial inter-class overlap indicated that physical scale alone was insufficient to accurately distinguish RBC morphotypes.

**Fig. 2.**
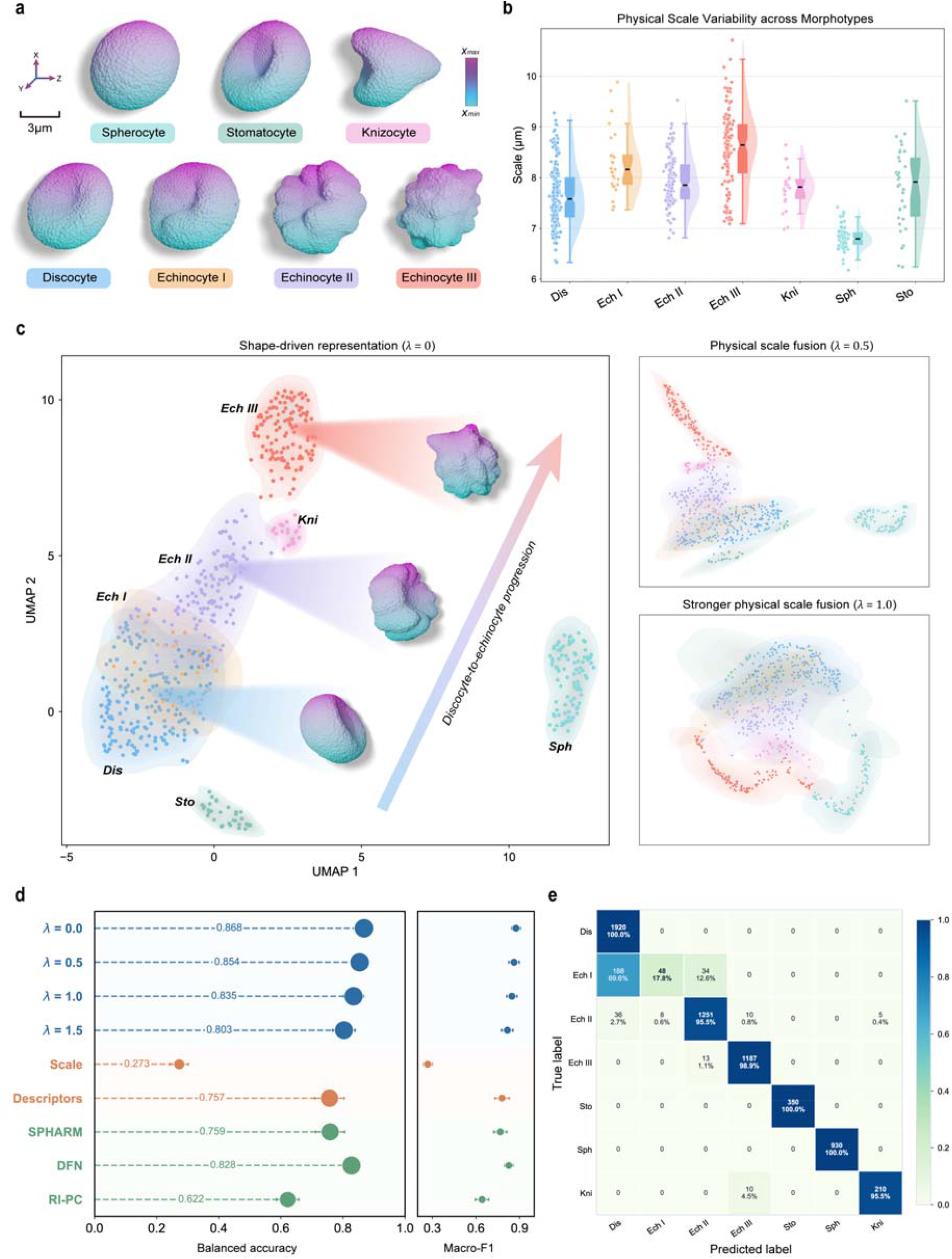
Shape-driven representations distinguish RBC morphotypes without fine-tuning. **a**, Representative 3D point clouds of seven RBC morphotypes, including discocyte, stomatocyte, spherocyte, knizocyte and echinocyte I–III^25^. Point clouds are colored by their *x* coordinate. Scale bar, 3 μm. **b**, Physical scale distributions across the seven RBC morphotypes. **c**, UMAP visualization of the shape-driven representation (*λ* = 0) and scale-fused representations (*λ* = 0.5 and 1.0). Each point represents one RBC and colors indicate morphotype. Representative point clouds illustrate the corresponding regions of the UMAP, and the arrow indicates the progression from discocytes to echinocytes. **d**, kNN classification performance across MorphCell representations with different physical scale fusion strengths (*λ* = 0, 0.5, 1.0 and 1.5), physical scale alone, 22 biophysical descriptors and three baseline representations, including spherical harmonics (SPHARM)^20^, Dynamic FoldingNet (DFN)^16^ and a rotation-invariant point cloud autoencoder (RI-PC)^15^. Performance was evaluated using balanced accuracy and macro-F1. Points and error bars indicate the mean and standard deviation, respectively. **e**, Confusion matrix for kNN classification using the MorphCell shape-driven representation.

We next asked whether MorphCell could capture morphotype-related variation in 3D shape independently of physical scale. We rescaled each point cloud to a common size before encoding, yielding a shape-driven representation (*λ* = 0), while retaining physical scale separately for controlled fusion (see Methods). Uniform manifold approximation and projection (UMAP)^26^ revealed a shape-dependent organization of these representations (Fig. 2c). Stomatocytes and spherocytes occupied relatively distinct regions, whereas discocytes and echinocytes I– III formed a continuous distribution. Echinocytes I were located predominantly between discocytes and echinocytes II, consistent with their intermediate degree of surface spiculation (Fig. 2a). Thus, without physical scale information or RBC-specific fine-tuning, MorphCell could identify RBC morphotypes based on their 3D shape.

We then examined how physical scale fusion affected the clustering. Increasing *λ* progressively altered the UMAP distributions (Fig. 2c and Supplementary Fig. 2a). At low fusion strength (*λ* = 0.5), separation among morphotypes was largely retained, whereas stronger fusion (*λ* = 1.0) produced greater overlap. Physical scale fusion therefore did not improve morphotype separation. To quantify morphotype discrimination, we compared the different representations using a kNN classification benchmark (Fig. 2d). The MorphCell shape-driven representation achieved the highest balanced accuracy and macro-F1 score across all representations evaluated. As *λ* increased, balanced accuracy declined from 86.8% to 80.3%, showing that stronger physical scale fusion weakened RBC morphotype discrimination. Physical scale alone only achieved a balanced accuracy of 27.3%, whereas the 22 biophysical descriptors achieved 75.7% (see Methods). We further compared the MorphCell shape-driven representation with three baseline representations, spherical harmonics (SPHARM)^20^, Dynamic FoldingNet (DFN)^16^ and a rotation-invariant point cloud autoencoder (RI-PC)^15^. MorphCell outperformed these baselines by 10.9%, 4.0% and 24.6%, respectively.

The confusion matrix for the shape-driven representation showed that classification errors were concentrated in echinocytes I (Fig. 2e). Misclassified echinocytes I were assigned primarily to discocytes and less frequently to echinocytes II. This classification result was consistent with the UMAP organization, in which echinocytes I lay between the smoother discocytes and the more spiculated echinocytes II (Fig. 2a,c).

These results demonstrate that pretrained MorphCell yielded transferable, shape-driven representations that distinguished RBC morphotypes without RBC-specific fine-tuning. In this dataset, 3D shape provided the primary discriminative information, whereas physical scale showed substantial within-morphotype variability, and its inclusion weakened rather than improved morphotype discrimination.

### MorphCell representations encode interpretable RBC shape variation

To interpret the RBC shape-driven representations learned by MorphCell, we related them to a set of biophysical descriptors spanning global structure and local surface detail (Fig. 3a–c; see Methods). Global descriptors included metrics such as volume, surface area, sphericity, solidity, elongation and flatness. Local descriptors quantified surface roughness, curvature and shape index across multiple spatial scales. We projected the shape-driven representations into principal component analysis (PCA) space and used Mantel tests to assess correlations between individual principal components and biophysical descriptors. Pairwise Spearman correlations were calculated among the descriptors (Fig. 3d; see Supplementary Note 3). PC1 explained 29.44% of the variance and was most strongly correlated with flatness, minor-axis length and the standard deviation of curvature at the largest spatial scale. By contrast, PC2 explained 12.84% of the variance and was correlated mainly with solidity, maximum roughness, and small- to medium-scale curvature and shape index statistics. These correlations linked PC1 primarily to global structural variation and PC2 to local surface variation.

**Fig. 3.**
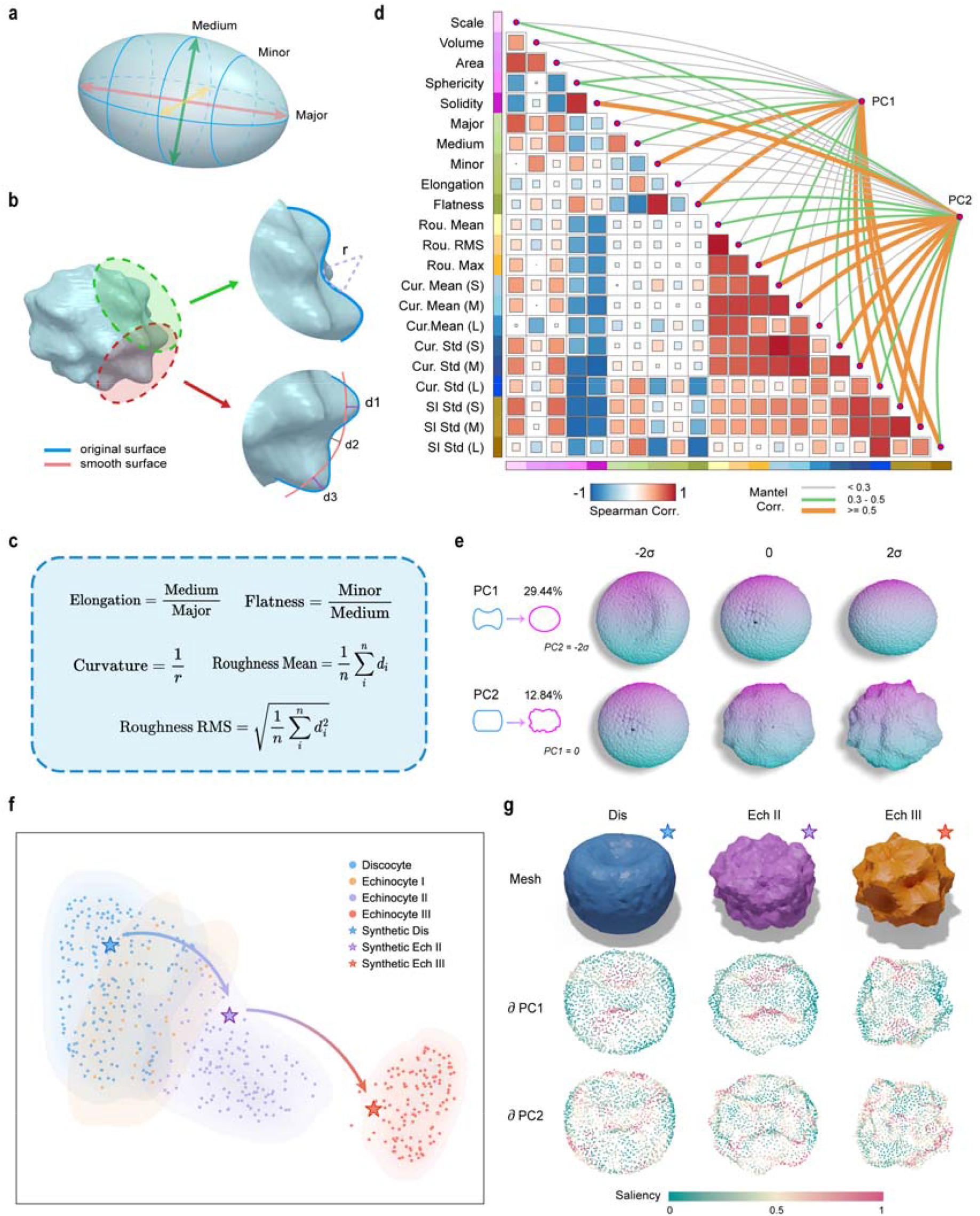
MorphCell representations encode interpretable RBC shape variation. **a**, Schematic of the PCA-derived major, medium and minor axes used to calculate elongation and flatness within the global structure descriptors. **b**, Schematics of local surface descriptors. Curvature is measured within a local neighborhood defined by radius *r*, whereas surface roughness is calculated from the distances between the original and smoothed surfaces. **c**, Representative formulas for elongation, flatness, curvature and surface roughness. **d**, Correlations between the principal components of the MorphCell shape-driven representations and biophysical descriptors. The matrix shows pairwise Spearman correlations among descriptors, and the links to PC1 and PC2 show Mantel correlations between individual principal components and descriptors. **e**, Interpolation along PC1 and PC2 followed by SQTD reconstruction. Reconstructed point clouds are shown at −2*σ*, 0 and 2*σ* along each principal component. Variation along PC1 changes the global contour from biconcave to near-spherical, whereas variation along PC2 increases the prominence of surface spicules. **f**, Projection of synthetic RBC samples onto a UMAP embedding constructed from real discocyte and echinocyte I– III samples. Stars denote the synthetic discocyte, echinocyte II and echinocyte III samples, which map to regions occupied by real cells with similar morphologies. **g**, Gradient saliency analysis of the synthetic RBC samples. PC1 saliency is concentrated in the polar depressions that form the biconcave contour, whereas PC2 saliency is concentrated on high-curvature surface spicules. Saliency values are normalized to the range 0-1.

To visualize the geometric variation represented by each principal component, we sampled along the principal component axes and reconstructed the corresponding point clouds using SQTD (Fig. 3e; see Supplementary Note 4). Along PC1, the reconstructed morphology transitioned continuously from a biconcave to a near-spherical contour while the surface remained relatively smooth. Along PC2, the overall contour changed less, whereas surface spicules became progressively more prominent. These reconstructions were consistent with the descriptor correlations identified by the Mantel tests (Fig. 3d).

We next examined whether the shape-driven representation responded to controlled geometric changes. We generated a sequence of synthetic RBCs ranging from a smooth discocyte to increasingly spiculated and spherical morphologies (see Supplementary Note 5).

When projected onto a UMAP embedding constructed from real discocyte and echinocyte I–III samples, the synthetic samples mapped to regions occupied by real cells with similar morphologies (Fig. 3f). Thus, the shape-driven representation tracked the controlled geometric changes introduced into the synthetic samples.

Finally, we applied gradient saliency analysis to the synthetic samples to identify the surface regions to which each principal component was most sensitive (Fig. 3g; see Supplementary Note 6). Saliency for PC1 was concentrated in the polar depressions that form the biconcave contour, whereas saliency for PC2 was concentrated on high-curvature surface spicules. These spatial patterns further supported the interpretation that PC1 primarily represented global contour, whereas PC2 primarily represented local surface detail.

### Shape and physical scale contribute differently to wheat root cell classification

To evaluate MorphCell in plant tissue, we applied it to a 3D X-ray microscopy (XRM) image stack of a wheat root. Cellpose-SAM^27^ was used for cell instance segmentation, enabling reconstruction of the root segment in 3D (Fig. 4a; see Methods). Wheat root cells exhibit distinct morphological characteristics across developmental zones and tissue regions. We therefore analyzed cell morphology across two axial developmental zones, the elongation and maturation zones, and five radial tissue regions, comprising the epidermis, cortex, endodermis, pericycle and stele (Fig. 4b,c).

**Fig. 4.**
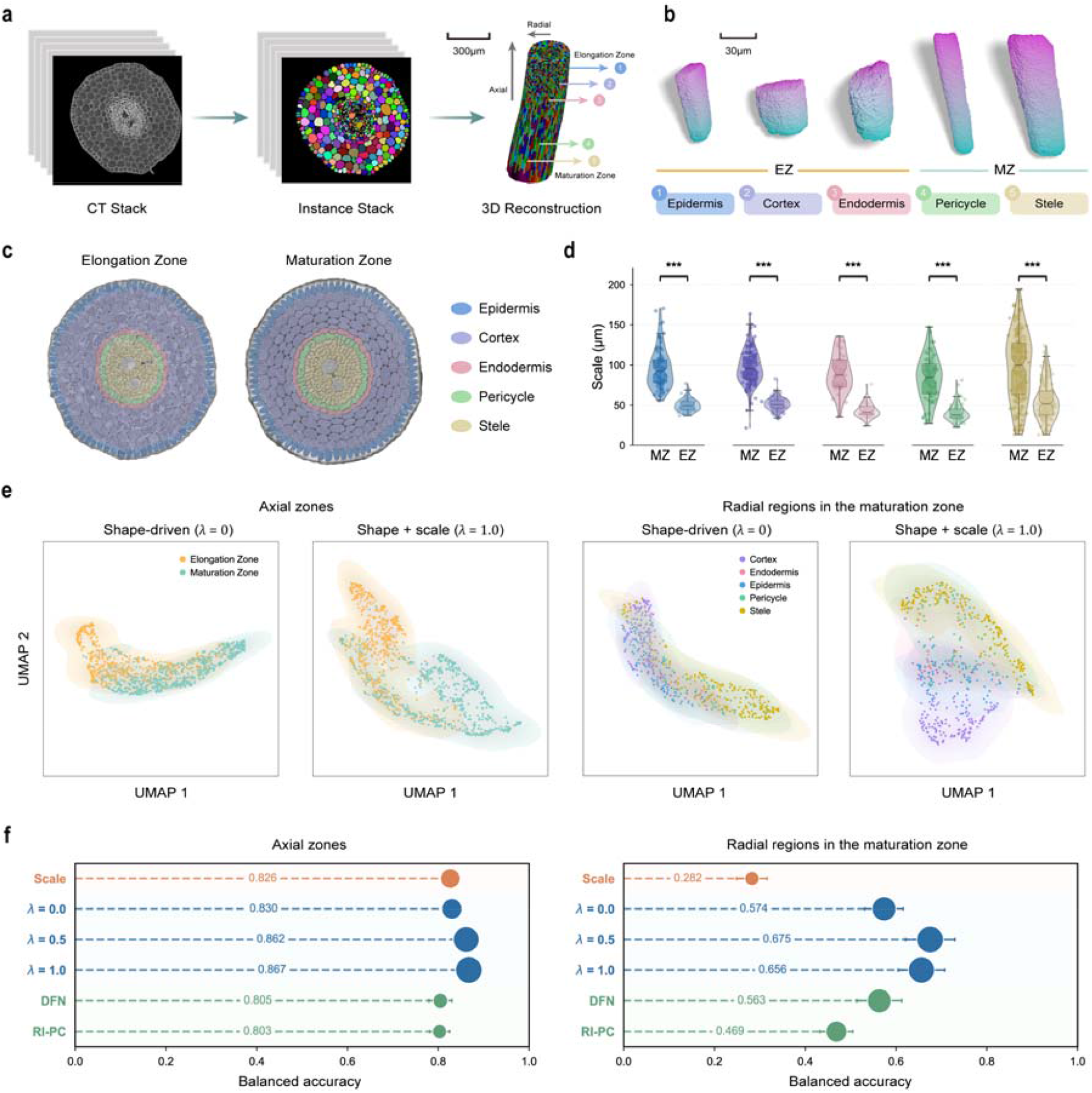
Shape and physical scale analysis of spatial organization in wheat root cells. **a**, Workflow for cell instance segmentation and 3D reconstruction from an X-ray microscopy (XRM) image stack of a wheat root. Cellpose-SAM segmentation produced an instance stack that was reconstructed into a 3D root segment with the axial and radial directions indicated. Scale bar, 300 μm. **b**, Representative 3D cell point clouds from the elongation zone (EZ) and maturation zone (MZ), illustrating the five radial tissue regions comprising the epidermis, cortex, endodermis, pericycle and stele. Scale bar, 30 μm. **c**, Cross-sectional views of the maturation and elongation zones, with the five radial tissue regions indicated. **d**, Physical scale distributions of MZ and EZ cells within each radial tissue region. MZ and EZ cells differed in physical scale across all five radial tissue regions (*P* < 0.001 for all comparisons). **e**, UMAP visualization of the shape-driven representation (*λ* = 0) and scale-fused representation (*λ* = 1.0) for the two axial zones and the five radial tissue regions within the maturation zone. **f**, kNN balanced accuracy for axial-zone classification and radial-region classification within the maturation zone using physical scale alone, MorphCell representations with different physical scale fusion strengths (*λ* = 0, 0.5 and 1.0), DFN and RI-PC.

We first examined physical scale and morphology representations across the axial zones and radial regions (Fig. 4d,e). Within each radial region, cells in the maturation zone were larger than those in the elongation zone (*P* < 0.001 for all comparisons). UMAP visualization of the shape-driven representation (*λ* = 0) partially separated elongation- and maturation-zone cells, although substantial overlap remained. Physical scale fusion (*λ* = 1.0) produced clearer separation between the two zones. For radial-region analysis, we focused on cells from the maturation zone. The shape-driven representation partially separated cells from different radial regions, with the clearest distinction between the cortex and stele. Physical scale fusion further separated some regions, whereas the epidermis, endodermis and pericycle remained substantially overlapping.

We then evaluated each representation using the kNN classification benchmark (Fig. 4f). For axial-zone classification, physical scale alone and the shape-driven representation yielded comparable balanced accuracies of 82.6% and 83.0%, respectively. Both exceeded those achieved by DFN (80.5%) and RI-PC (80.3%). Physical scale fusion further improved balanced accuracy to 86.2% at *λ* = 0.5 and 86.7% at *λ* = 1.0. For radial-region classification within the maturation zone, physical scale alone yielded a balanced accuracy of only 28.2%, whereas the shape-driven representation achieved 57.4%, exceeding DFN (56.3%) and RI-PC (46.9%). Physical scale fusion further improved balanced accuracy, reaching the highest value of 67.5% at *λ* = 0.5. These results indicate that both 3D shape and physical scale were important for classifying axial zones and radial tissue regions, but their relative contributions differed between the two biological tasks.

### Joint cell and nucleus representations distinguish colon epithelial cell states

To evaluate the ability of MorphCell to characterize disease-associated 3D cellular morphology, we applied it to a vEM dataset of cancerous and non-cancerous epithelial cells from one patient with colon cancer (Fig. 5a). We used U-Net-CEM500K^28^ and SAM2-assisted interactive segmentation^29^ to extract nucleus and cell masks, respectively (Fig. 5b d; see Methods). The SAM2-derived cell masks were subsequently evaluated and corrected by experts. The final dataset contained 70 colon epithelial cells, including 47 cancerous cells and 23 non-cancerous cells, each with paired 3D cell and nucleus structures.

**Fig. 5.**
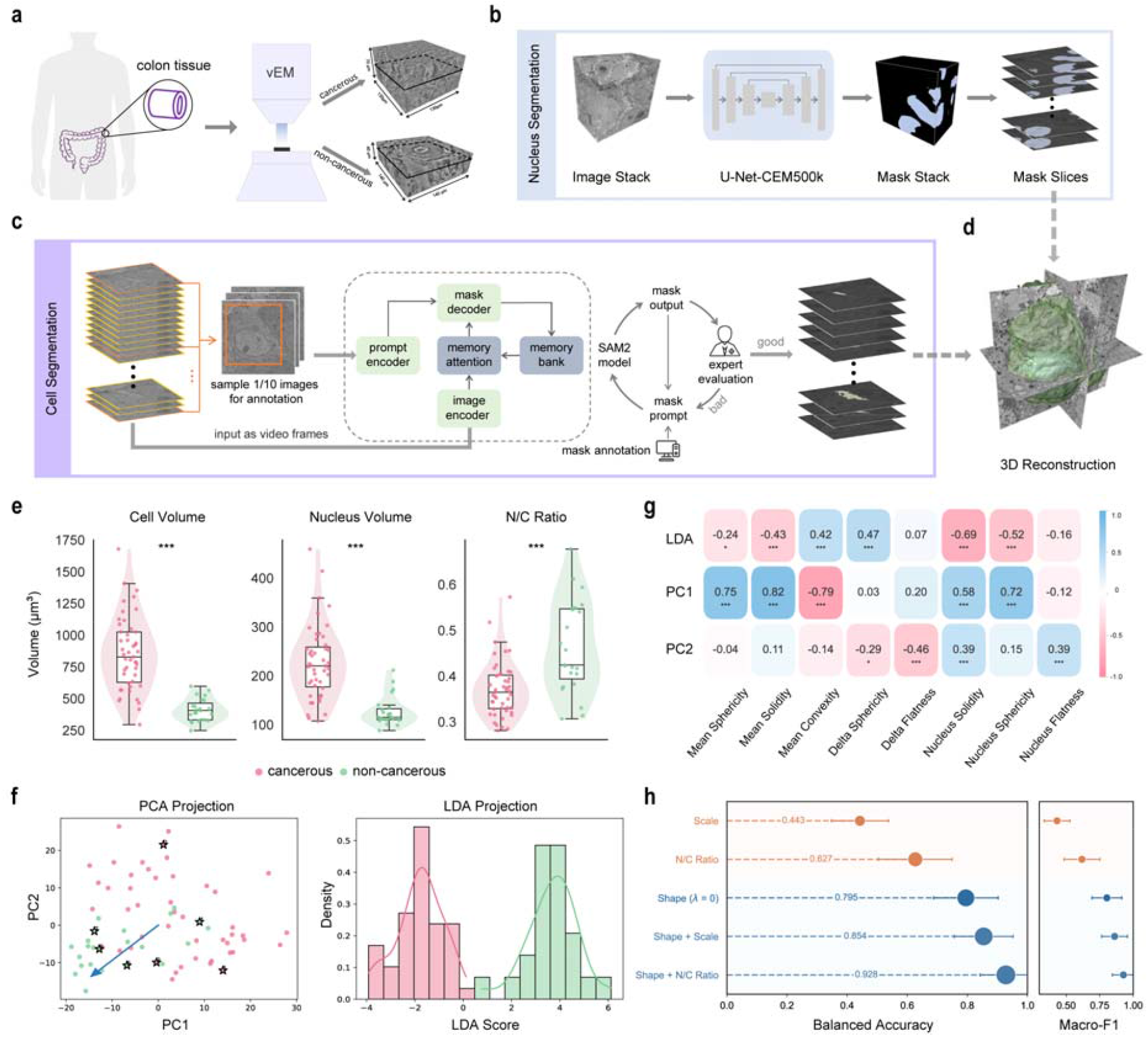
Joint analysis of cell and nucleus morphology in colon epithelial cells. **a**, Overview of the vEM dataset from histologically confirmed tumor and adjacent non-tumor tissues of a patient with colon cancer. **b**, Nucleus segmentation workflow using U-Net-CEM500K^28^ to generate nucleus mask stacks. **c**, Cell segmentation workflow using SAM2-assisted interactive segmentation^29^. Annotated sections were used as prompts, propagated through the image stack and iteratively corrected following expert evaluation. **d**, 3D reconstruction of paired cell and nucleus structures. **e**, Comparisons of cell volume, nuclear volume and the nuclear-to-cytoplasmic volume ratio (N/C ratio) between cancerous and non-cancerous cells. Statistical significance is indicated above each comparison. **f**, PCA and LDA projections of the joint cell–nucleus representations. **g**, Correlations between the LDA score, principal components and biophysical descriptors. **h**, kNN classification performance for cancerous and non-cancerous cells using physical scale, N/C ratio, the shape-driven representation (*λ* = 0), and the shape-driven representation fused with either physical scale or N/C ratio (*λ* = 0.5 for both).

We first compared cell volume, nuclear volume and the nuclear-to-cytoplasmic volume ratio (N/C ratio) between cancerous and non-cancerous cells (Fig. 5e). Cancerous cells had larger cell and nuclear volumes than non-cancerous cells, consistent with the nuclear enlargement commonly associated with malignant epithelial morphology^30^. However, the increase in cell volume exceeded that in nuclear volume, resulting in a lower N/C ratio in cancerous cells. We therefore asked whether these volume differences were accompanied by differences in physical scale. Cell physical scale, nucleus physical scale and the nucleus-to-cell scale ratio showed no significant differences between the two groups (Supplementary Fig. 3a). Thus, the differences in cell and nuclear volume were not reflected in corresponding differences in physical scale.

We next extracted shape-driven representations separately from the cell and nuclear surfaces. The two representations were concatenated to form a joint cell–nucleus representation, which was analyzed using PCA and linear discriminant analysis (LDA)^20^ (Fig. 5f; see Supplementary Note 2). PCA showed partial separation between cancerous and non-cancerous cells. LDA further separated the two groups along a supervised discriminant axis. These results show that the joint cell – nucleus representation captured morphological differences between the two groups beyond physical scale. To interpret these representation differences, we correlated the LDA score and principal components with biophysical descriptors (Fig. 5g; see Supplementary Note 3). The LDA score was strongly associated with nuclear solidity, nuclear sphericity and the cell–nucleus difference in sphericity. PC1 was mainly associated with mean convexity and mean solidity. PC2 showed weaker correlations, primarily with the cell–nucleus difference in flatness and nuclear flatness.

Finally, we used a kNN classification benchmark to assess the contributions of physical scale, N/C ratio and the shape-driven representation (Fig. 5h and Supplementary Fig. 3b). Physical scale showed the weakest discrimination, whereas the N/C ratio was more informative but remained less discriminative than the shape-driven representation. Combining the shape-driven representation with either measurement improved classification, with N/C ratio fusion achieving the highest balanced accuracy of 92.8%. Thus, joint cell–nucleus shape provided the primary discriminative information, while the N/C ratio contributed complementary volume information.

## Discussion

3D cell morphology contains information that is not fully captured by conventional size measurements or 2D imaging techniques. In this study, we developed MorphCell, a point-cloud representation learning framework for 3D cellular morphology. Without biological task-specific training, MorphCell learns transferable shape-driven representations and reconstructs complete surfaces from individual representations while retaining physical scale as a separate measurement. Applied to RBCs, wheat root cells and paired cell–nucleus structures from colon epithelium, MorphCell captured variation in both global contour and local surface detail. The reconstruction branch links variation in the learned representations to the underlying 3D geometry, while separate scale measurements allow shape and physical scale to be evaluated independently or in combination. These mechanisms of MorphCell provide a unified representation space for deciphering 3D cellular morphology across biological systems and imaging modalities.

The transferability of MorphCell across biological datasets was supported by its cross-view reconstruction pretraining on diverse 3D surface geometries from ShapeNetCore V2^31^. During pretraining, independent cropping and rotation generated two geometrically related views in separate coordinate systems, requiring the model to infer structural relationships between spatially distinct surface regions rather than simply reproduce the observed input. Consistent with this design, our ablation experiment showed that cross-view reconstruction pretraining improved downstream performance across the evaluated biological transfer tasks compared with direct reconstruction pretraining (Supplementary Note 1 and Supplementary Table 2). MorphCell also performed better than the self-reconstruction-based DFN^16^ and RI-PC^15^ models in the RBC morphotype and wheat root cell classification tasks. These results suggest that cross-view reconstruction can provide an effective self-supervised strategy for learning reusable geometric representations from unlabeled 3D surface data.

Cellular morphology encompasses both physical scale and geometric shape, which may carry distinct biological information. MorphCell explicitly separates the shape-driven representation from physical scale, allowing their individual and combined contributions to classification to be quantitatively evaluated. Across the biological applications examined here, the relative contributions differed according to the classification task. In RBCs, the stomatocyte-discocyte-echinocyte sequence reflects the mechanical properties of the membrane and its associated cytoskeleton, together with constraints imposed by membrane area and cell volume^32,33^. Such mechanical constraints can generate substantial changes in cell shape without equivalent changes in maximum cell diameter, consistent with our finding that shape-driven representations were more informative than physical scale for RBC morphotype classification. In wheat roots, axial development involves both an increase in cell size and directionally biased expansion governed by cell wall mechanics^34,35^. The former contributes to changes in physical scale, whereas the latter alters cell proportions and 3D geometry, providing a biological basis for the complementary information carried by shape and physical scale in axial-zone classification. For radial-region classification within the maturation zone, shape-driven representations provided the primary discriminative information, while incorporating physical scale increased classification accuracy. In colon epithelial cells, the joint cell–nucleus representation captured morphological differences between cancerous and non-cancerous cells beyond physical scale. Correlations with biophysical descriptors of cell and nuclear morphology helped interpret these representation differences. The N/C ratio provided additional discriminative information when combined with the shape-driven representation. These analyses suggest that MorphCell can reveal biologically interpretable morphological variation that is not fully captured by conventional size measurements alone.

In summary, MorphCell provides a unified framework for 3D cellular morphology analysis. Future work could expand the pretraining datasets to include more diverse geometries, which may improve the representation of irregular and branched biological forms. Additionally, extending this framework to explicitly model the spatial relationships among cellular and subcellular structures could enable the analysis of integrated cellular architecture.

## Methods

### Datasets

#### ShapeNetCore V2

MorphCell was pretrained on ShapeNetCore V2^31^, a widely used benchmark for 3D point cloud processing in computer vision. The dataset contains 52,470 mesh models from 55 everyday object categories, including furniture, vehicles and aircraft. We uniformly sampled 2,048 surface points from each mesh as the model input.

#### Red blood cells

For zero-shot morphological analysis, we used a published 3D confocal microscopy dataset of RBCs from healthy donors and patients with hereditary spherocytosis^25^. Cells were imaged using a Nikon Eclipse Ti inverted microscope equipped with a CSU-W1 spinning-disk confocal head (Yokogawa) and a 60× oil-immersion objective with a numerical aperture of 1.4. The image stacks had voxel dimensions of 110 × 110 × 300 nm. From the available RBC morphologies, we retained seven morphotypes for analysis: discocytes, stomatocytes, spherocytes, knizocytes and echinocytes I–III.

#### Wheat root cells

The dataset comprised a 3D image stack of a wheat root acquired using a Zeiss Xradia 610 Versa X-ray microscope. The stack contained 1,997 slices with an isotropic voxel size of 0.7 µm. It covered the elongation and maturation zones, with five radial tissue regions including the epidermis, cortex, endodermis, pericycle and stele.

#### Colon epithelial cells

Colon epithelial tissue samples were obtained from one patient with colon cancer. Cancerous and non-cancerous cells were derived from histologically confirmed tumor and adjacent non-tumor tissues, respectively. Samples were imaged using a Helios G3 focused ion beam scanning electron microscope (FIB-SEM) to generate vEM stacks. The final analysis included 70 epithelial cells, comprising 47 cancerous and 23 non-cancerous cells. Each image stack contained the cell surface and its corresponding nucleus, enabling joint analysis of their 3D morphology. Voxel dimensions were 5.61523 × 5.61523 × 50 nm for cancerous cells and 7.324 × 7.324 × 50 nm for non-cancerous cells.

#### IntrA

IntrA is an open-access dataset of 3D intracranial vasculature reconstructed from time-of-flight magnetic resonance angiography (TOF-MRA) scans of human brains^24^. It comprises 103 complete cerebrovascular models, 1,909 automatically generated vessel segments, including 1,694 healthy and 215 aneurysm-containing segments, and 116 aneurysm segments manually annotated by medical experts. We selected one vessel segment for reconstruction analysis.

### Segmentation

#### Wheat root cells

Individual wheat root cells were segmented using a fine-tuned Cellpose-SAM model^27^. We fine-tuned the model using manually annotated slices from the XY, XZ and YZ planes of the wheat root stack. During inference, 256 × 256 sliding windows were applied to slices in each orientation to predict 2D flow fields. The three sets of flow predictions were combined into a 3D flow field across the complete stack. Gradient tracking followed this field and grouped voxels converging at the same location into individual cell masks. Noisy and boundary-truncated instances were removed.

#### Colon epithelial cells

For each colon epithelial cell, the cell surface and nucleus were segmented separately. Nuclei were segmented automatically using U-Net-CEM500K^28^, which comprised a ResNet50 encoder with 23 million parameters and a U-Net decoder with 9 million parameters. The encoder had been pretrained on CEM500K using MoCoV2 unsupervised contrastive learning. CEM500K contains 500,000 curated and deduplicated 2D cellular electron microscopy images. We fine-tuned U-Net-CEM500K on 100 manually annotated nucleus images.

Cell surfaces were segmented using SAM2-assisted interactive segmentation^29^. Each vEM stack was treated as a video, with its constituent slices serving as frames. We manually annotated bounding boxes on one of every ten slices and supplied them as prompts to SAM2. During inference, the image encoder processed each frame, while the memory module retained information from the annotated slices and propagated it through the remaining stack to generate a sequence of cell masks.

Experts reviewed the propagated masks and provided additional correction prompts for slices with discontinuous or inaccurate cell contours. SAM2 then updated the masks for the current and subsequent slices. This evaluation and correction process was repeated until the masks were spatially continuous and suitable for quantitative analysis.

### Point cloud preprocessing

Point cloud preprocessing began from either grayscale image stacks or segmented cell masks. RBC data were provided as grayscale stacks, which required filtering and binary segmentation before mask processing. By contrast, colon epithelial and wheat root cells were already available as segmented masks.

RBC image stacks were smoothed using a 3D Gaussian filter with a standard deviation of *σ* = 1.0. Otsu’s maximum between-class variance threshold and Li’s minimum cross-entropy threshold were calculated for each stack. Their arithmetic mean was used as the global threshold to reduce sensitivity to uneven fluorescence and background noise.

After RBC segmentation, all three datasets entered the same mask processing workflow. We applied morphological hole filling to remove internal cavities caused by local signal loss and discarded components smaller than 100 voxels. We then used 3D connected component labeling to retain only the largest component. A continuous triangular mesh was extracted from each mask using marching cubes at an isovalue of 0.5. We corrected the surface normals to maintain a consistent mesh orientation.

We used farthest point sampling (FPS)^36^ to select 2,048 approximately uniform points from the mesh vertices. Principal component analysis (PCA) was then applied to standardize point cloud orientation. Its eigenvectors defined a rotation matrix that aligned the longest principal axis with the x axis of a common Cartesian coordinate system.

### Model architecture

MorphCell comprised a point cloud encoder^23^ for cross-view reconstruction and a Spherical-Query Transformer Decoder (SQTD) for self-reconstruction.

#### Cross-view reconstruction

During pretraining, each aligned point cloud was independently cropped twice to generate two local views. The crop ratio for each view was sampled uniformly between 0.6 and 1.0, and the two views were allowed to overlap. Each view was normalized to the unit sphere and randomly rotated, placing the views in separate coordinate systems.

We divided each view into local patches. FPS selected 64 patch centers, and the 32 nearest points to each center formed a patch. A patch embedding layer converted each patch into a 384-dimensional token. The shared 12-layer Transformer encoder^37^ then encoded these tokens into a view-specific representation.

Reconstruction was performed in both directions, with one view serving as the source and the other as the target. View-Relative Positional Embedding (VRPE) provided the spatial queries required to recover the target patches from the source representation. Let *c*_*i*_ ∈ ℝ^3^ denote the center coordinate of target patch *i*, and let *r* ∈ ℝ^3^ denote the global relative displacement between the two views. We concatenated these terms into a six-dimensional positional vector:

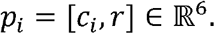

The positional embedding function *ϕ*(·) mapped this vector to a high-dimensional VRPE query:

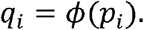

We used *q*_*i*_ as the cross-attention query and the source representation *F*_*s*_ as both the keys and values. The output for target patch *i* was

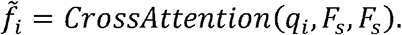

The cross-attention outputs 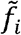 were passed through a four-layer Transformer decoder and then linearly projected into 3D coordinates to reconstruct the target patches. Reversing the source and target views produced the reconstruction in the opposite direction. For downstream analyses, complete point clouds were grouped into patches and passed through the encoder without view cropping or random rotation.

#### Spherical self-reconstruction

Cross-view reconstruction could not recover a complete point cloud from a single representation because its VRPE queries required target patch centers and the relative displacement between two views. We therefore developed SQTD to reconstruct the complete morphology without this view-specific spatial information.

SQTD began with 2,048 uniformly sampled points on a Fibonacci sphere as a fixed template. To reduce excessive smoothing when using 3D coordinates directly as queries, we applied Fourier embedding to each template coordinate:

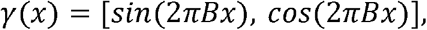

where *x* denotes a template coordinate and *B* is the Fourier projection matrix. The values *γ*(*x*) served as queries in a four-layer Transformer decoder, whereas the encoder representation of the reconstruction target provided the keys and values. A multilayer perceptron then predicted a 3D displacement *Δ*_*i*_ for each template point *x*_*i*_. The reconstructed coordinate was therefore 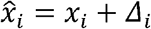.

We initialized the weights and biases of the final linear layer to zero, so the initial decoder output remained close to the spherical template. During training, the predicted displacements progressively deformed this template into the target morphology.

### Training

MorphCell was pretrained on ShapeNetCore V2 using self-supervised learning. We used the dataset only as a source of diverse 3D surface geometry and did not use its category labels. We therefore pooled the official training and test splits, yielding 52,470 mesh models. All downstream biological datasets were excluded from pretraining.

Before training, each mesh was resampled to 2,048 points by FPS with an expansion ratio of 1.1. We applied online augmentation through random rotation around the y axis and Gaussian coordinate jittering (*σ* = 0.01, clipped to ±0.05). Point clouds were also anisotropically scaled between 0.8 and 1.2 and randomly translated within ±0.1;.

Cross-view reconstruction and spherical self-reconstruction were optimized jointly. The total loss was the weighted sum of their reconstruction losses:

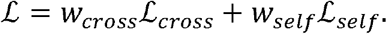

Both objectives used bidirectional Chamfer distance^38^. Given a predicted point cloud *P* and a target point cloud *Q*, the loss was

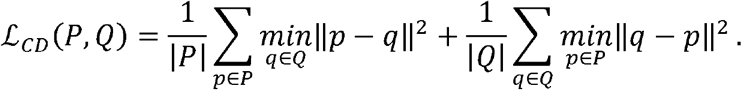

For cross-view reconstruction, the loss was calculated separately for each target patch and summed across both reconstruction directions. For spherical self-reconstruction, the loss was calculated over all points in each view and summed across the two views. Cross-view reconstruction was assigned a weight of *w*_*cross*_ = 1.0, whereas spherical self-reconstruction served as an auxiliary objective with *w_self_* = 0.5.

We optimized the model using AdamW^39^ with a weight decay of 0.05. The learning rate was linearly increased from 10^−6^ to 5×10^−4^ during a 10-epoch warm-up, followed by 290 epochs of cosine annealing^40^ to 10^−6^. Training ran for 300 epochs with a batch size of 16 and a random seed of 42. The model was trained on a single NVIDIA A800 GPU for approximately two days.

### Physical scale fusion

To construct scale-fused representations, we first applied PCA to the MorphCell shape-driven representations. We retained components explaining 80% of the cumulative variance, yielding the reduced matrix *F_pca_*.

Physical scale *S* was calculated from the original point cloud coordinates. It was defined as twice the maximum distance from any point to the point cloud centroid. This value corresponded to the maximum physical diameter.

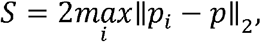

where *p*_*i*_ denotes point *i* and *p* is the point cloud centroid. Physical scale values were standardized by z-score transformation.

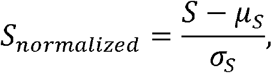

where *μ*_*S*_ and *σ*_*S*_ denote the mean and standard deviation of physical scale, respectively. We used variation along PC1 as the reference for physical scale fusion. Let *σ*_*PC*1_ denote the standard deviation of PC1 scores. Each scale-fused representation was constructed as

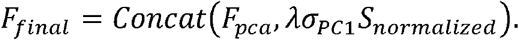

The hyperparameter *λ* controlled the contribution of physical scale relative to variation along PC1. When *λ* =0, physical scale was excluded, yielding the shape-driven representation. At *λ* = 0.5, 1.0 and 1.5, the appended physical scale values had standard deviations equal to 0.5, 1.0 and 1.5 times *σ* _*PC*1_, respectively.

### kNN classification benchmark

We used a k-nearest-neighbor (kNN) classification benchmark to compare representations. The evaluated inputs comprised MorphCell representation variants, physical scale, biophysical descriptors and baseline representations. MorphCell variants were constructed with *λ* ∈ {0,0.5,1.0,1.5}. The shape-driven representation corresponded to *λ* = 0, whereas *λ* >0 produced scale-fused representations. Physical scale alone was evaluated using *S*_*normalized*_. For the biophysical baseline, the 22 descriptors other than physical scale were standardized within each training fold. Baseline representations comprised rotation-invariant spherical harmonic descriptors (SPHARM)^20^, Dynamic FoldingNet (DFN)^16^ and a rotation-invariant point cloud autoencoder (RI-PC)^15^. All inputs were evaluated using a kNN classifier with *k=5* and Euclidean distance. Neighbor contributions were weighted by the inverse of their distance.

Performance was evaluated using repeated stratified fivefold cross-validation with ten repeats, yielding 50 train-test splits. A random seed of 42 was used to generate the splits. The classifier was fitted on each training split and evaluated on the corresponding test split. Performance was summarized as the mean and standard deviation across the 50 test splits. Predictions from all test splits were accumulated to construct the confusion matrix.

The primary metrics were balanced accuracy and macro-F1. Balanced accuracy was defined as the arithmetic mean of class-specific recall:

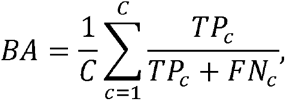

where *C* is the number of classes, and *TP*_*C*_ and *FN*_*C*_ denote true positives and false negatives for class *c*. Macro-F1 was the unweighted mean of the class-specific F1 scores:

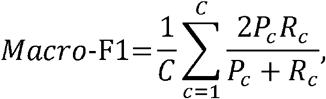

where *P*_*C*_ and *R*_*C*_ are the precision and recall for class *c*, respectively.

### Biophysical descriptors

We calculated biophysical descriptors from triangular surface meshes to interpret MorphCell representations. These descriptors quantified global structure and local surface detail (Supplementary Table 1).

#### Global structure

Global descriptors comprised physical scale, volume *v*, surface area, sphericity, solidity, convexity, principal axis lengths, elongation and flatness. Volume and surface area were calculated from each closed triangular mesh. Sphericity quantified similarity to an ideal sphere. Solidity was calculated as the ratio of mesh volume to convex hull volume. Convexity was calculated as the ratio of mesh surface area to convex hull surface area.

We applied PCA to the mesh vertices to obtain the major, medium and minor axis lengths, denoted *L*_*maj*_, *L*_*med*_ and *L*_*min*_, respectively. Elongation was defined as *L*_*med*_/*L*_*maj*_, and flatness as *L*_*min*_/*L*_*med*_.

For colon epithelial cells, global descriptors were calculated separately for the cell and nucleus. For each descriptor, the mean value was calculated by averaging the corresponding cell and nucleus values. The delta value was calculated by subtracting the nucleus value from the cell value.

#### Local detail

We applied ten iterations of Laplacian smoothing to each mesh to generate a reference surface with reduced high-frequency detail. Local roughness *d*_*i*_ was defined as the Euclidean distance between vertex i in the original and smoothed meshes. The mean, root mean square and maximum of *d*_*i*_ were used as roughness descriptors.

We estimated curvature at each vertex using discrete curvature measures within a spherical neighborhood of radius *r*. Mean curvature *H* was calculated from dihedral angle contributions and normalized by the mesh surface area within the neighborhood. Gaussian curvature K was calculated from vertex angle deficits and normalized by the same area. Curvature was evaluated at small, medium and large spatial scales using radii of *r* = 3.0, 8.0 and 15.0 voxels, respectively. At each scale, mean curvature magnitude was calculated as the mean of |*H*|, avoiding cancellation between positive and negative values. The standard deviation of *H* quantified spatial variation in surface bending. Gaussian curvature was not included as a separate descriptor but was combined with *H* to calculate the shape index^41^.

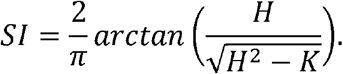

The shape index ranges from −1 to 1 and distinguishes local geometries such as cup-like, saddle-like and ridge-like surfaces. Its standard deviation was calculated at each spatial scale to quantify spatial variation in local surface geometry.

## Resource availability

### Lead contact

Further information and requests for resources and reagents should be directed to, and will be fulfilled by, the lead contact, Xuping Feng.

### Materials availability

This study did not generate new materials.

### Data and code availability

The MorphCell datasets, including 3D image volumes and processed point clouds, are available in Science Data Bank at https://doi.org/10.57760/sciencedb.0132h. The data-processing code is provided at https://github.com/Visytudz/MorphCell-Datasets. The MorphCell source code is available at https://github.com/Visytudz/MorphCell.

## Acknowledgements

We thank the Bio-ultrastructure Analysis Laboratory at the Analysis Center of Agrobiology and Environmental Sciences, Zhejiang University, for assistance with cell imaging.

## Author information

**College of Biosystems Engineering and Food Science, Zhejiang University, Hangzhou 310058, China**.

Guowei Zhang, Runzhou Cao, Shan Xu, Zeyu Yu, Yong He, Xuping Feng

**School of Artificial Intelligence and Information Engineering, Zhejiang University of Science and Technology, Hangzhou, China**.

Guan Wang

**Center of Cryo-Electron Microscopy, Zhejiang University School of Medicine, Hangzhou 310058, China**.

Jiansheng Guo

**Westlake Laboratory of Life Sciences and Biomedicine, Hangzhou, China**.

Yiyan Zheng

**The Rural Development Academy & Agricultural Experiment Station, Zhejiang University, Hangzhou 310058, China**.

Xuping Feng

## Author contributions

Guowei Zhang, Guan Wang and Runzhou Cao contributed equally to this work. Guowei Zhang, Xuping Feng, Runzhou Cao and Shan Xu conceived the study and designed the experiments. Guowei Zhang, Guan Wang, Jiansheng Guo, Shan Xu and Zeyu Yu performed data acquisition and curation. Guowei Zhang and Xuping Feng wrote the manuscript. Yong He and Yiyan Zheng reviewed and edited the manuscript.

## References

1. Seal, S. et al. Cell Painting: a decade of discovery and innovation in cellular imaging. Nat. Methods 22, 254–268 (2025).

2. Ramezani, M. et al. A genome-wide atlas of human cell morphology. Nat. Methods 22, 621–633 (2025).

3. Guan, G. et al. Cell lineage-resolved embryonic morphological map reveals signaling associated with cell fate and size asymmetry. Nat. Commun. 16, 3700 (2025).

4. Villeneuve, C. et al. Mechanical forces across compartments coordinate cell shape and fate transitions to generate tissue architecture. Nat. Cell Biol. 26, 207–218 (2024).

5. Lovegrove, H. E. et al. Interphase cell morphology defines the mode, symmetry, and outcome of mitosis. Science 388, eadu9628 (2025).

6. Xie, S., Zhang, S., de Medeiros, G., Liberali, P. & Skotheim, J. M. The G1/S transition in mammalian stem cells in vivo is autonomously regulated by cell size. Nat. Commun. 16, 9071 (2025).

7. Phillip, J. M., Han, K.-S., Chen, W.-C., Wirtz, D. & Wu, P.-H. A robust unsupervised machine-learning method to quantify the morphological heterogeneity of cells and nuclei. Nat Protoc 16, 754–774 (2021).

8. Theis, S., Mendieta-Serrano, M. A., Chapa-y-Lazo, B., Chen, J. & Saunders, T. E. CellMet: Extracting 3D shape and topology metrics from confluent cells within tissues. PLOS Computational Biology 21, e1013260 (2025).

9. Chandrasekaran, S. N., Ceulemans, H., Boyd, J. D. & Carpenter, A. E. Image-based profiling for drug discovery: due for a machine-learning upgrade? Nat Rev Drug Discov 20, 145–159 (2021).

10. Zinchenko, V., Hugger, J., Uhlmann, V., Arendt, D. & Kreshuk, A. MorphoFeatures for unsupervised exploration of cell types, tissues, and organs in volume electron microscopy. eLife 12, e80918 (2023).

11. Daetwyler, S. & Fiolka, R. P. Light-sheets and smart microscopy, an exciting future is dawning. Commun. Biol. 6, 502 (2023).

12. Sakdinawat, A. & Attwood, D. Nanoscale X-ray imaging. Nature Photon. 4, 840–848 (2010).

13. Peddie, C. J. et al. Volume electron microscopy. Nat. Rev. Methods Primers 2, 51 (2022).

14. Driscoll, M. K. et al. Robust and automated detection of subcellular morphological motifs in 3D microscopy images. Nat. Methods 16, 1037–1044 (2019).

15. Vasan, R. et al. Interpretable representation learning for 3D multi-piece intracellular structures using point clouds. Nat. Methods 22, 1531–1544 (2025).

16. De Vries, M. et al. Geometric deep learning and multiple-instance learning for 3D cell-shape profiling. Cell Systems 16, 101229 (2025).

17. Ong, H. T. et al. Digitalized organoids: integrated pipeline for high-speed 3D analysis of organoid structures using multilevel segmentation and cellular topology. Nat. Methods 22, 1343–1354 (2025).

18. Ducroz, C., Olivo-Marin, J.-C. & Dufour, A. Spherical Harmonics based extraction and annotation of cell shape in 3D time-lapse microscopy sequences. Annu. Int. Conf. IEEE Eng. Med. Biol. Soc. 2011, 6619–6622 (2011).

19. Medyukhina, A. et al. Dynamic spherical harmonics approach for shape classification of migrating cells. Sci. Rep. 10, 6072 (2020).

20. Viana, M. P. et al. Integrated intracellular organization and its variations in human iPS cells. Nature 613, 345–354 (2023).

21. Burgess, J. et al. Orientation-invariant autoencoders learn robust representations for shape profiling of cells and organelles. Nat. Commun. 15, 1022 (2024).

22. Murthy, R. S. et al. Generalizable morphological profiling of cells by interpretable unsupervised learning. Nat. Commun. 16, 11465 (2025).

23. Zhang, X., Zhang, S. & Yan, J. Towards more diverse and challenging pre-training for point cloud learning: Self-supervised cross reconstruction with decoupled views. in Proceedings of the IEEE/CVF international conference on computer vision (ICCV) 28696–28706 (2025).

24. Yang, X., Xia, D., Kin, T. & Igarashi, T. IntrA: 3D intracranial aneurysm dataset for deep learning. in Proceedings of the IEEE/CVF conference on computer vision and pattern recognition (CVPR) (2020).

25. Simionato, G. et al. Red blood cell phenotyping from 3D confocal images using artificial neural networks. PLoS Comput. Biol. 17, e1008934 (2021).

26. McInnes, L., Healy, J. & Melville, J. UMAP: Uniform Manifold Approximation and Projection for Dimension Reduction. Preprint at 10.48550/arXiv.1802.03426 (2020).

27. Pachitariu, M., Rariden, M. & Stringer, C. Cellpose-SAM: superhuman generalization for cellular segmentation. Preprint at 10.1101/2025.04.28.651001 (2025).

28. Conrad, R. & Narayan, K. CEM500K, a large-scale heterogeneous unlabeled cellular electron microscopy image dataset for deep learning. eLife 10, e65894 (2021).

29. Ravi, N. et al. SAM 2: Segment anything in images and videos. in International Conference on Learning Representations (ICLR) (2025).

30. Li, Y. et al. Nuclear Softness Promotes the Metastatic Potential of Large-Nucleated Colorectal Cancer Cells via the ErbB4-Akt1-Lamin A/C Signaling Pathway. International Journal of Biological Sciences 20, 2748–2762 (2024).

31. Chang, A. X. et al. ShapeNet: An Information-Rich 3D Model Repository. Preprint at 10.48550/arXiv.1512.03012 (2015).

32. Khairy, K., Foo, J. & Howard, J. Shapes of Red Blood Cells: Comparison of 3D Confocal Images with the Bilayer-Couple Model. Cel. Mol. Bioeng. 1, 173–181 (2008).

33. Geekiyanage, N. M. et al. A coarse-grained red blood cell membrane model to study stomatocyte-discocyte-echinocyte morphologies. PLOS ONE 14, e0215447 (2019).

34. Verbelen, J.-P., Cnodder, T. D., Le, J., Vissenberg, K. & Baluška, F. The Root Apex of Arabidopsis thaliana Consists of Four Distinct Zones of Growth Activities: Meristematic Zone, Transition Zone, Fast Elongation Zone and Growth Terminating Zone. Plant Signaling & Behavior 1, 296–304 (2006).

35. Cosgrove, D. J. Structure and growth of plant cell walls. Nat. Rev. Mol. Cell Biol. 25, 340–358 (2024).

36. Qi, C. R., Yi, L., Su, H. & Guibas, L. J. PointNet++: deep hierarchical feature learning on point sets in a metric space. in Proceedings of the 31st international conference on neural information processing systems 5105–5114 (Curran Associates Inc., Long Beach, California, USA, 2017).

37. Vaswani, A. et al. Attention is all you need. in Proceedings of the 31st international conference on neural information processing systems (eds Guyon, I. et al.) vol. 30 (Curran Associates, Inc., Long Beach, California, USA, 2017).

38. Fan, H., Su, H. & Guibas, L. A Point Set Generation Network for 3D Object Reconstruction from a Single Image. in 2017 IEEE Conference on Computer Vision and Pattern Recognition (CVPR) 2463–2471 (IEEE, Honolulu, HI, 2017).

39. Loshchilov, I. & Hutter, F. Decoupled weight decay regularization. in International Conference on Learning Representations (ICLR) (2019).

40. Loshchilov, I. & Hutter, F. SGDR: Stochastic gradient descent with warm restarts. in International Conference on Learning Representations (ICLR) (2017).

41. Koenderink, J. J. & van Doorn, A. J. Surface shape and curvature scales. Image and Vision Computing 10, 557–564 (1992).

